# Lymphatic metastases have more diverse roots than distant metastases

**DOI:** 10.1101/828913

**Authors:** Johannes G. Reiter, Shriya Nagpal, Kamila Naxerova

## Abstract

Both lymphatic and distant metastases arise through cancer cell migration and colonization of ectopic sites. Nonetheless, the two metastasis types are associated with significantly different clinical outcomes, suggesting that distinct biological mechanisms may drive their formation. Here we show fundamental differences in the seeding patterns of lymphatic and distant metastases. Analyzing the reconstructed phylogenies of human colorectal cancers, we find that distant metastases typically are monophyletic, originating from one common ancestor. Lymphatic metastases, in contrast, are almost exclusively polyphyletic and can be seeded from many primary tumor regions. We develop a rigorous mathematical framework for quantifying the phylogenetic diversity of metastases while accounting for differential lesion sampling among patients. Our results indicate that a smaller fraction of primary tumor cells gives rise to distant metastases than lymphatic metastases. Thus, the two metastasis types exhibit profoundly distinct phylogenetic traits, indicating that different evolutionary mechanisms may drive their formation and influence their clinical behavior.

## Main text

In most cancers, metastasis to distant organs confers a considerably worse prognosis than spread to locoregional lymph nodes. For example, 5-year survival for colorectal cancers that have metastasized to local lymph nodes or the pericolonic fat (stage III) is 53-90% but drops to 12% for patients with spread to distant organs (stage IV)^1^. The survival difference for patients with locoregional and distant disease is similar for other tumor types, such as breast cancer and melanoma^2,3^.

The formation of both lymphatic and distant metastases depends on cancer cell migration and colonization of foreign microenvironments^4^. Given that both types of metastasis require similar cellular abilities^5^ and indicate the presence of a potentially lethal cell type capable of ectopic growth, it is worth asking why clinical outcomes of stage IV patients differ so markedly from those of stage III patients.

The simplest explanation is that distant metastases often affect vital organs such as the liver and the lungs and therefore lead to accelerated death. However, locoregional recurrence may be equally dangerous in some cancer types. For example, autopsy studies have shown that local recurrence was the cause of death in approximately 50% of colorectal cancer patients^6^, highlighting the importance of locoregional disease control. Similarly, in pancreatic cancer, local recurrence has been estimated to be responsible for approximately 30% of deaths^7^.

Are lymphatic metastases perhaps easier to remove than distant metastases? 5-year survival for colorectal cancer patients with resectable liver metastases is 25-44%^8^, well above average for stage IV disease, suggesting that surgical management of metastases can make a difference. Yet, clinically, resection of affected lymph nodes is not a high priority in colorectal cancer. Nodes are primarily resected for staging and not for therapeutic purposes^9^. Pre-operative imaging of mesenteric lymph nodes is challenging^10^ and lymph node harvest practices vary by institution^11^. Therefore, affected nodes probably stay behind in a fraction of patients. In rectal cancer, clinical trials have shown that extended lateral pelvic lymph node dissection did not improve survival^12,13^, echoing similar findings in breast cancer^14^ and melanoma^15^. Collectively, these data suggest that “left-behind” positive nodes do not necessarily lead to local recurrence and call into question the idea that relative ease of surgical management is the reason for the survival difference between patients with lymphatic and distant metastases.

Finally, distant and lymphatic metastases may represent fundamentally different disease forms that are driven by distinct biology and dissemination mechanisms. To date, no systematic comparative studies have investigated the evolutionary features of lymphatic and distant metastases in humans. Here, we show that lymph node metastases in colorectal cancer are a phylogenetically more diverse group than distant metastases. Genetic heterogeneity among lymph node metastases mirrors the genetic diversity of the primary tumor. Phylogenetic analyses show that lymphatic metastases intermingle with primary tumor regions on the evolutionary tree, indicating that in stage III patients, many if not all primary tumor regions are capable of seeding lymph node metastases. In stark contrast, distant metastases are a homogeneous, monophyletic group that tends to be the terminal branch of the phylogenetic tree. Their distinctive phylogenetic features indicate that lymphatic and distant metastases arise from cancer cells with different biological properties.

To investigate the evolution of lymphatic and distant metastases, we took advantage of a recently published collection of colorectal cancer phylogenies^16^. From this study, we selected all patients (n=18) with multiple primary tumor regions (range 2-10) and/or lymph node and/or distant metastases (range 2-10). These data formed the basis of our analysis (see Supplementary Table 1 for detailed patient information). Importantly, this cohort was exhaustively sampled, and a majority of resected metastases of sufficient size and purity were included, minimizing sampling bias^16^. Phylogenies were reconstructed based on small insertions and deletions in hypermutable polyguanine tracts, a method that produces rich mutation data and robust trees^17^. We had previously used this patient cohort to ascertain that most liver metastases originate in the primary tumor and do not share a common subclonal origin with lymph node metastases^16^. Here, we analyzed the evolutionary trees from a fundamentally different perspective, asking whether lymphatic and distant metastases *as a group* consistently display distinct phylogenetic features.

Evaluating patient trees (**Fig. 1a** and Supplementary Figures 1-3), we noticed a recurring pattern. Lymph node metastases and primary tumor samples typically diverged, often in alternating succession, from the tree trunk, while distant lesions usually had one common ancestor and tended to form the terminal branch of the tree (Supplementary Fig. 4). Given the consistency of this observation, we sought to formalize it. First, to avoid sampling bias, we reduced the data set to one sample per lymphatic and distant metastasis. That is, in cases where multiple biopsies were taken from the same metastasis, we randomly removed all but one sample, such that each metastasis was represented by only one biopsy in the final data set (see Supplementary Figs. 1-3 for all phylogenies). Then, we determined the fraction of patients in whom all anatomically distinct distant metastases had one common ancestor and grouped together in a monophyletic clade that did not include any primary tumors or lymphatic metastases. We found that in 67% of patients, distant metastases were part of such a clade. In contrast, lymphatic metastases formed a monophyletic group in only 10% of patients (**Fig. 1b**, p = 0.036, two-tailed Fisher’s exact test). A slightly altered classification approach in which we considered distant and lymphatic metastases a monophyletic group if the clade contained all metastases but no primary tumor samples (allowing for the other metastasis type to be part of the branch) gave similar results, with 20% and 83% of patients having one common ancestor for all lymphatic and distant metastases, respectively (p = 0.035, two-tailed Fisher’s exact test, Supplementary Fig. 5). Note that the classification into monophyletic and polyphyletic groups is unrelated to our previously described common and distinct origin categories, which reflect whether lymphatic and distant metastases have a common subclonal origin^16^. For example, all phylogenetic trees in Fig. 1a show polyphyletic lymph node metastases and monophyletic distant metastases, although C45 and C66 belong to the distinct origins category, while C36 shows common origin of lymphatic and distant metastases^16^.

**Fig. 1:**
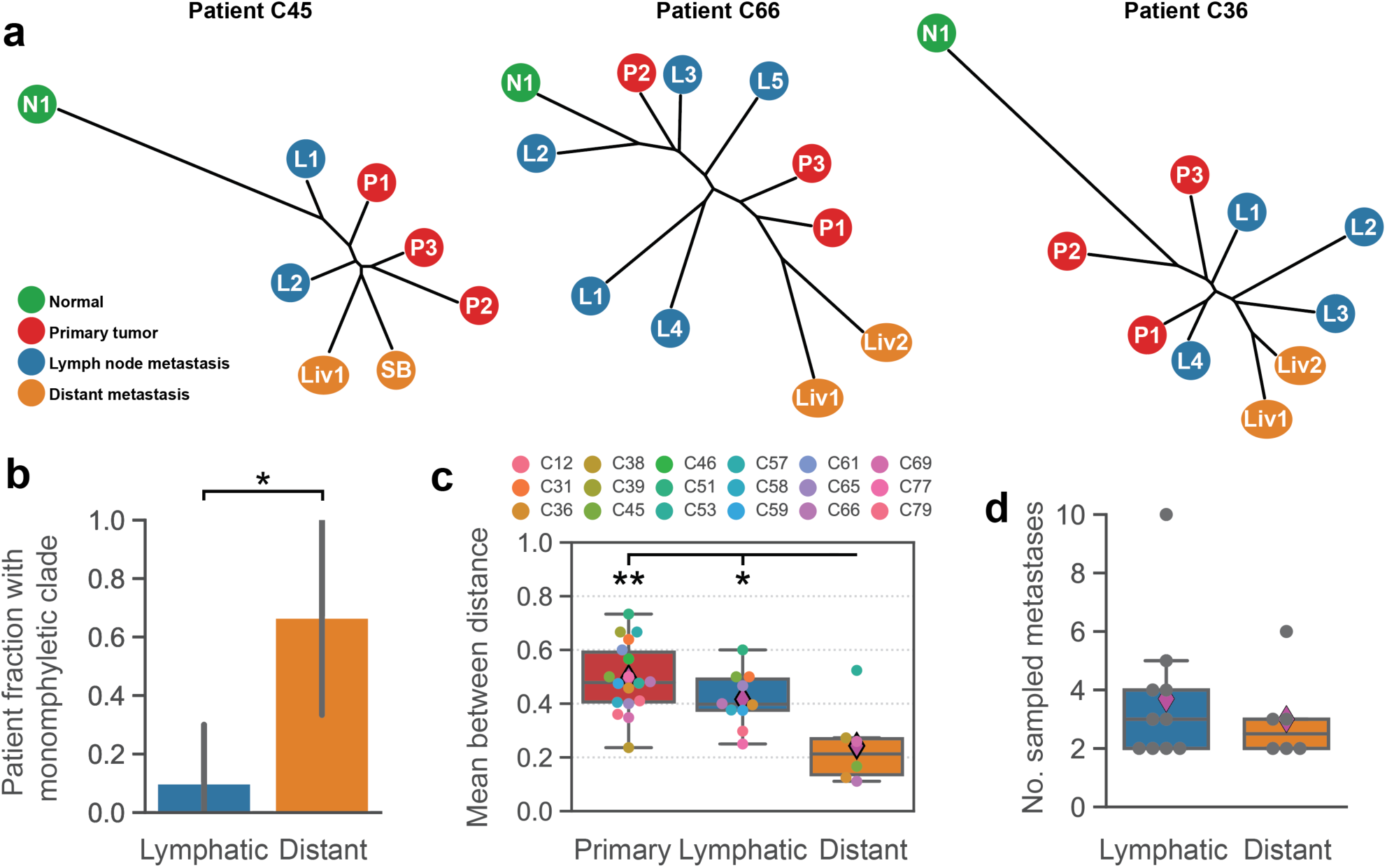
Distant but not lymphatic metastases form monophyletic clades in most patients. **a |** Phylogenetic trees of colorectal cancer patients C45, C66, C36, adapted from Naxerova et al.^16^. Distant metastases arise from a common ancestor in all cases. Liv, liver metastasis; SB, small bowel metastasis. **b |** All distant metastases formed a monophyletic clade in 67% (4/6) of patients. All lymphatic metastases formed a monophyletic group in 10% (1/10) of patients (p = 0.036, two-tailed Fisher’s exact test). The black bars denote 90% confidence intervals. **c |** The normalized mean number of internal phylogenetic nodes that separated a pair of distinct distant metastases was significantly lower than the mean for primary tumor samples (0.24 vs 0.5) or lymphatic metastases (0.24 vs 0.42), respectively. No statistically significant difference was observed between the mean distances of primary tumor samples and lymphatic metastases (p=0.11, two-tailed Mann-Whitney test). Center line, median; box limits, upper and lower quartiles; points, outliers. Magenta diamonds illustrate the mean in each group. **d |** No statistically significant difference was observed between the number of lymphatic and distant metastases samples (mean of 3.7 vs 3; p=0.54, two-tailed t-test). * *P < 0.05*; ** *P < 0.01*.

We further explored the high phylogenetic homogeneity of distant metastases (**Fig. 1b**) by calculating, for every patient, the mean phylogenetic distance (number of internal nodes) separating different primary tumor regions and distinct lymphatic and distant metastases. The distances were not significantly different for primary tumor regions and lymphatic metastases (mean distances of 0.50 vs 0.42) but significantly lower for distant metastases (mean distance of 0.24, p=8.4e-3 and p=0.045, two-tailed Mann-Whitney tests), confirming the relative homogeneity of this group (**Fig. 1c**).

We wondered whether differential sampling of lymph node and distant metastases may have affected the results. For example, if ten lymphatic but only two distant metastases are included in a phylogeny, it is far more likely that all distant metastases will have one common ancestor by chance. We did not observe a significant difference between the number of lymphatic and distant metastases in our data set, but the mean and variance were slightly higher in the lymph node metastasis group (mean 3.7 vs 3.0 metastases, p=0.54, Student’s t-test, **Fig. 1d**). Additionally, the number of primary tumor regions sampled in each case further affects the odds of finding monophyletic metastasis groups by chance alone. To account for the different number of lesions sampled in each patient, we developed a mathematical framework that allowed us to quantify the likelihood of common origin for any given phylogeny. We define *m* as the number of metastasis samples under investigation (either lymphatic or distant), and *k* as the number of all other tumor samples in the phylogeny (**Supplementary Methods**). We calculate a *root diversity score* (RDS) defined by the probability that at least *l* out of *m* metastases form a common clade in a tree with *n* = *k* + *m* samples (Supplementary Table 2). In other words, the root diversity score denotes the probability that a tree with an equally or more extreme clustering of metastases occurs by chance alone. For example, in subject C36 (**Fig. 1a**), the root diversity score for distant metastasis is 0.067, as the likelihood that two distant metastases (m=2) will cluster by chance in a phylogeny with n=9 samples is 6.7%. The power to detect non-random clustering of metastases increases with the number of samples *n* in a phylogeny (**Fig. 2a**).

**Fig. 2.**
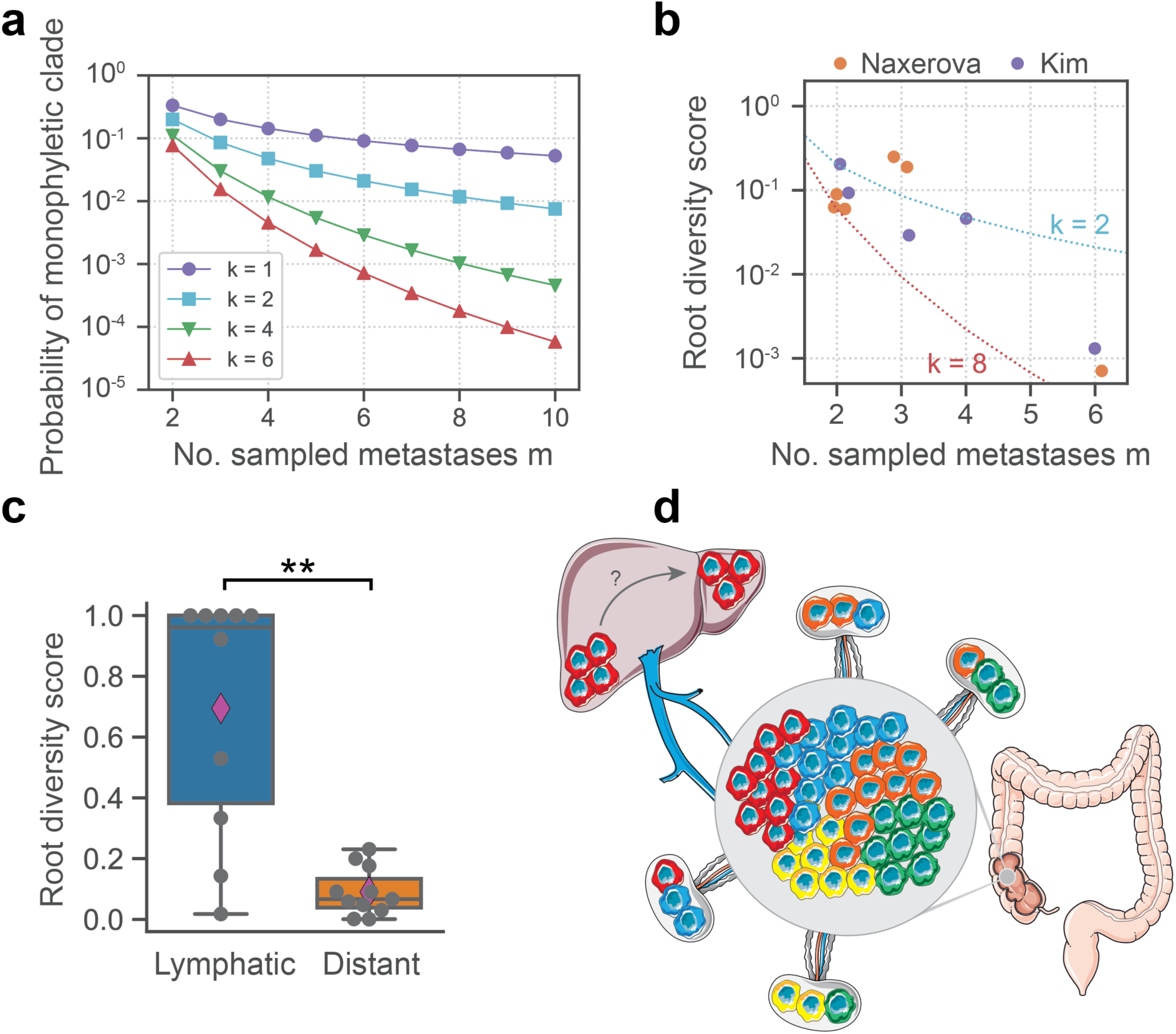
Distant but not lymphatic metastases exhibit a very low root diversity score. **a |** The probability of observing a monophyletic clade of all sampled metastases *m* by chance decreases with increasing *m* and increasing number of other cancer samples *k*. **b |** In both cohorts, the root diversity score decreases as the power to observe a low score increases with the number of sampled distant metastases. k ranges between 2 and 8 in both cohorts. **c |** The root diversity score was significantly lower for distant metastases than lymphatic metastases (0.09 vs 0.65; p=0.0026; two-tailed Mann-Whitney test). Center line, median; box limits, upper and lower quartiles; points, outliers. Magenta diamonds illustrate the mean in each group. **d |** Summary schematic showing that lymphatic metastases can be seeded from many primary tumor regions and mirror the heterogeneity of the primary tumor, while distant metastases are typically formed by one clone, either due to selection or intra-organ metastasis.

We used the root diversity score to quantify the homogeneity of distant metastases in our cohort. We found that after accounting for the number of other samples (k) in the phylogenies, indeed the root diversity score was generally very low (**Fig. 2b**), even for phylogenies where distant metastases did not form a monophyletic clade. To validate the low root diversity of distant metastases in an independent cohort, we searched the literature for colorectal cancer phylogenies with at least two primary tumor samples and multiple anatomically distinct distant lesions. We found one appropriate study comprising five patients with a total of 17 liver metastases^18^. We calculated the root diversity scores for distant metastases for all five patients (trees are shown in Supplementary Fig. 6) and found the smallest possible root diversity score in every case (**Fig. 2b**), independently confirming our observation that distant metastases tend to be monophyletic. In 8 out of 11 patients with multiple distant lesions in the combined two cohorts, the likelihood that metastases would cluster to the observed degree by chance alone was below 10% (Supplementary Table 2). Furthermore, combining all root diversity scores, we calculated a combined p-value of 4.5e10^−7^ for the entire patient population. This p-value corresponds to the likelihood that distant metastases would cluster to the observed degree by chance. Thus, we find strong evidence for distant metastasis homogeneity both within individual phylogenies and across the whole patient cohort.

Returning to our original question, we next applied the root diversity score to lymphatic and distant metastases in a comparative analysis. The results showed highly significant differences in root diversity between the two metastasis types (mean diversity score of 0.69 vs 0.090; p=2.6e10^−3^, two-tailed Mann-Whitney test), confirming that lymphatic metastases are far more likely to be polyphyletic than distant metastases (**Fig. 2c**), even after accounting for differential sampling in a mathematically rigorous fashion.

We wondered whether these differences might be due to treatment effects. Treatment did not affect the majority of patients in the combined two cohorts, as 16 out of 23 cases (70%) had synchronous metastasis. In these cases, all primary and metastatic lesions were resected at the same time. Seven patients had metachronous metastasis and received treatment in the time interval between the resection of the primary tumor and associated lymphatic metastases and the resection of distant metastases. Two of these did not have multiple distant metastases (C65, C39) and therefore were not included in **Fig. 2c**. Only five patients with multiple distant metastases had metachronous metastasis and treatment in the interval between resections (C66, C36, C69, CRC2 and CRC5). In one case (C69), some distant metastases (Liv1, Liv2) were resected along with the primary tumor and the lymph nodes, and others (Liv3) 6 months later, after a chemotherapy regimen. All distant metastases still clustered together in this case (Supplementary Fig. 2), arguing against an effect of the treatment on the inferred phylogeny. Nonetheless, we recalculated the root diversity score after excluding *all* treated patients (C66, C36, C69, CRC2 and CRC5) from the analysis and found that the results remained highly significant (7.0e10^−3^, Supplementary Fig. 7).

In summary, our results indicate that in colorectal cancer, lymphatic and distant metastases are phylogenetically distinct groups. Lymph node metastases are polyphyletic, mirror the heterogeneity of the primary tumor and are furthermore polyclonal, according to a recent report^19^. These observations suggest the absence of strong selection during the formation of lymph node metastases: many cells from the primary tumor appear capable of migrating to and thriving in lymph nodes. Distant metastases, in contrast, typically have one common ancestor and form a monophyletic group (**Fig. 2d**).

Multiple explanations for the high phylogenetic similarity of distant metastases exist. First, metastases may have given rise to each other^20–22^. Most lesions in our data set were liver metastases and could have formed through intra-hepatic seeding. Standing on its own, we consider this explanation relatively unlikely, as many phylogenetically similar metastases (e.g. C69, C36, CRC3, CRC4) presented in different liver segments, which are independent functional units with separate vascular systems. Furthermore, the two patients in our cohort who had metastases in different organs (C45 and C38) still showed monophyletic origin of these lesions.

Second, it is possible that distant metastasis represents a specific selective bottleneck and thereby, in contrast to lymphatic metastasis, selects for a particular subpopulation. The ability to enter and exit the blood stream^23^, travel longer distances^24^, or survive in organ-specific microenvironments^25^ may represent such a bottleneck. This possibility is further supported by a recent study which showed that distant metastases in different cancer types were more often monophyletic than expected by chance^26^. The existence of an (epi-) genetically defined metastatic clone has been strongly debated over the years^27^. Our results motivate a continued search for the molecular traits of this clone. It will furthermore be important to determine whether metastasis to different organs selects for different lineages^25^, a question that cannot be conclusively answered with our liver-centric data set.

Most importantly, our data show that lymphatic metastases evolve by fundamentally different rules than distant metastases in colorectal cancer. Lymphatic metastases’ phylogenetic features reflect the relative absence of strong selective pressures, and no specialized clone appears to be necessary for their formation, potentially explaining their more benign clinical implications.

## Methods

### Root diversity score

The root diversity score (RDS) denotes the probability that in a cancer phylogeny with n tumor samples at least l out of m metastases samples form a single clade. We generalized Edwards’ and Cavalli-Sforza’s approach to calculate the number of distinct phylogenies with a given number of samples in which at least l of m metastases samples form a monophyletic group^28,29^ (**Supplementary Methods**). To obtain the probability that such a phylogeny would evolve by chance, we divide this number of phylogenies by the total number of phylogenies with n tumor samples (see Equation S2 in **Supplementary Methods**). All RDS values are provided in Supplementary Table 2.

### Code availability

The source code to calculate the root diversity score as well as to produce various figure panels is available as jupyter notebook at http://www.github.com/johannesreiter/rootdiversity. (The code will be released upon publication; for review please see supplementary files). The notebooks are implemented in Python 3.6. All required input data is contained in Supplementary Tables 1 and 2.

### Data availability

Results are based on previously published data and inferred cancer phylogenies. Original raw polyguanine profiling data, and phylogenetic trees can be downloaded from datadryad.org (http://dx.doi.org/10.5061/dryad.vv53d). Original whole-exome sequencing data of Kim et al.^18^ was deposited to the Sequence Read Archive (SRA) at the NCBI under the project ID of PRJNA271316. All figures have associated raw data.

## Acknowledgments

We would like to thank Christina Curtis and Martin Nowak for helpful discussions. This work was support by grants from the NIH (R37CA225655) and AACR (561314) to K.N. and from NCI (K99CA229991) to J.G.R.

## Author contributions

K.N. and J.G.R. designed the study. All authors performed the research and wrote the manuscript.

## References

1. Survival Rates for Colorectal Cancer, by Stage. American Cancer Society Available at: https://www.cancer.org/cancer/colon-rectal-cancer/detection-diagnosis-staging/survival-rates.html#references. (Accessed: 11th December 2018)

2. Survival Rates for Melanoma Skin Cancer, by Stage. American Cancer Society Available at: https://www.cancer.org/cancer/melanoma-skin-cancer/detection-diagnosis-staging/survival-rates-for-melanoma-skin-cancer-by-stage.html. (Accessed: 11th December 2018)

3. Breast Cancer Survival Rates & Statistics. American Cancer Society Available at: https://www.cancer.org/cancer/breast-cancer/understanding-a-breast-cancer-diagnosis/breast-cancer-survival-rates.html. (Accessed: 11th December 2018)

4. Massagué, J. & Obenauf, A. C. Metastatic colonization by circulating tumour cells. Nature 529, 298–306 (2016).

5. Wong, S. Y. & Hynes, R. O. Lymphatic or Hematogenous Dissemination: How Does a Metastatic Tumor Cell Decide? Cell Cycle 5, 812–817 (2006).

6. Welch, J. P. & Donaldson, G. A. The clinical correlation of an autopsy study of recurrent colorectal cancer. Ann. Surg. 189, 496–502 (1979).

7. Iacobuzio-Donahue, C. A. et al. DPC4 gene status of the primary carcinoma correlates with patterns of failure in patients with pancreatic cancer. J. Clin. Oncol. 27, 1806–13 (2009).

8. Garden, O. J. et al. Guidelines for resection of colorectal cancer liver metastases. Gut 55 Suppl 3, iii1–8 (2006).

9. Chen, S. L. & Bilchik, A. J. Resecting Lymph Nodes in Colon Cancer: More than a Staging Operation? Ann. Surg. Oncol. 14, 2175–2176 (2007).

10. O’Dwyer, S. T., Haboubi, N. Y., Johnson, J. S. & Gardy, R. Detection of lymph node metastases in colorectal carcinoma. Colorectal Dis. 3, 288–94 (2001).

11. McDonald, J. R., Renehan, A. G., O’Dwyer, S. T. & Haboubi, N. Y. Lymph node harvest in colon and rectal cancer: current considerations. World J Gastrointest Surg 2012; 4:9–19.

12. Georgiou, P. et al. Extended lymphadenectomy versus conventional surgery for rectal cancer: a meta-analysis. Lancet. Oncol. 10, 1053–62 (2009).

13. Georgiou, P. A., Mohammed Ali, S., Brown, G., Rasheed, S. & Tekkis, P. P. Extended lymphadenectomy for locally advanced and recurrent rectal cancer. Int. J. Colorectal Dis. 32, 333–340 (2017).

14. Giuliano, A. E. et al. Axillary Dissection vs No Axillary Dissection in Women With Invasive Breast Cancer and Sentinel Node Metastasis. JAMA 305, 569 (2011).

15. Faries, M. B. et al. Completion Dissection or Observation for Sentinel-Node Metastasis in Melanoma. N. Engl. J. Med. 376, 2211–2222 (2017).

16. Naxerova, K. et al. Origins of lymphatic and distant metastases in human colorectal cancer. Science 357, 55–60 (2017).

17. Naxerova, K. et al. Hypermutable DNA chronicles the evolution of human colon cancer. Proc. Natl. Acad. Sci. U. S. A. 111, E1889–98 (2014).

18. Kim, T.-M. et al. Subclonal Genomic Architectures of Primary and Metastatic Colorectal Cancer Based on Intratumoral Genetic Heterogeneity. Clin. Cancer Res. 21, 4461–72 (2015).

19. Ulintz, P. J., Greenson, J. K., Wu, R., Fearon, E. R. & Hardiman, K. M. Biology of Human Tumors Lymph Node Metastases in Colon Cancer Are Polyclonal. Clin Cancer Res 24, (2017).

20. Gundem, G. et al. The evolutionary history of lethal metastatic prostate cancer. Nature 520, 353–357 (2015).

21. McPherson, A. et al. Divergent modes of clonal spread and intraperitoneal mixing in high-grade serous ovarian cancer. Nat. Genet. 48, 758–767 (2016).

22. El-Kebir, M., Satas, G. & Raphael, B. J. Inferring parsimonious migration histories for metastatic cancers. Nat. Genet. 50, 718–726 (2018).

23. Valastyan, S. & Weinberg, R. A. Tumor metastasis: molecular insights and evolving paradigms. Cell 147, 275–92 (2011).

24. Ryser, M. D., Min, B.-H., Siegmund, K. D. & Shibata, D. Spatial mutation patterns as markers of early colorectal tumor cell mobility. Proc. Natl. Acad. Sci. U. S. A. 115, 5774–5779 (2018).

25. Obenauf, A. C. & Massagué, J. Surviving at a distance: organ specific metastasis. Trends in cancer 1, 76–91 (2015).

26. Zhao, Z.-M. et al. Early and multiple origins of metastatic lineages within primary tumors. Proc. Natl. Acad. Sci. U. S. A. 113, 2140–5 (2016).

27. Vanharanta, S. & Massagué, J. Origins of Metastatic Traits. Cancer Cell 24, 410–421 (2013).

28. Edwards, A. W. F., Cavalli-Sforza, L.. The reconstruction of evolution. Heredity (Edinb). 18, 104–105 (1963).

29. Felsenstein, J. Inferring phytogenies. Sinauer Assoc. (2003). doi:10.1086/383584

