## Supplementary Materials for "Lymphatic metastases have more diverse roots than distant metastases"

[*]Correspondence to:

### Supplementary Figures

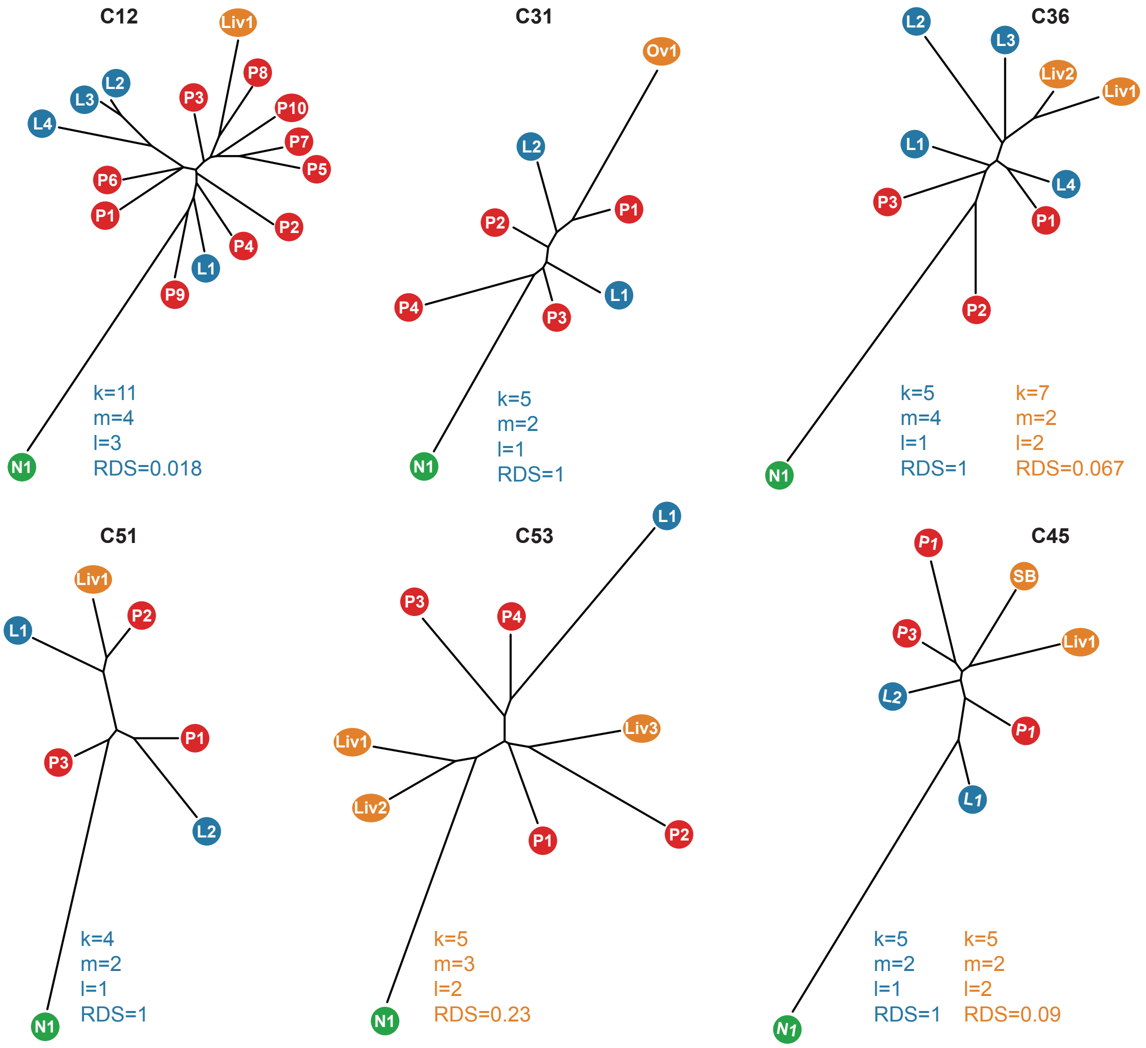


**Supplementary Fig. 1: Phylogenetic trees of colorectal cancer patients C12, C31, C36, C51, C53, and C45.** Trees adapted from Naxerova et al.[16]. N, normal tissue; P, primary tumor; L, lymph node metastasis; Liv, liver metastasis; Ov, ovarian metastasis, SB, small bowel; RDS, root diversity score. The RDS of lymphatic metastases is denoted in blue. The RDS of distant metastases is denoted in orange.

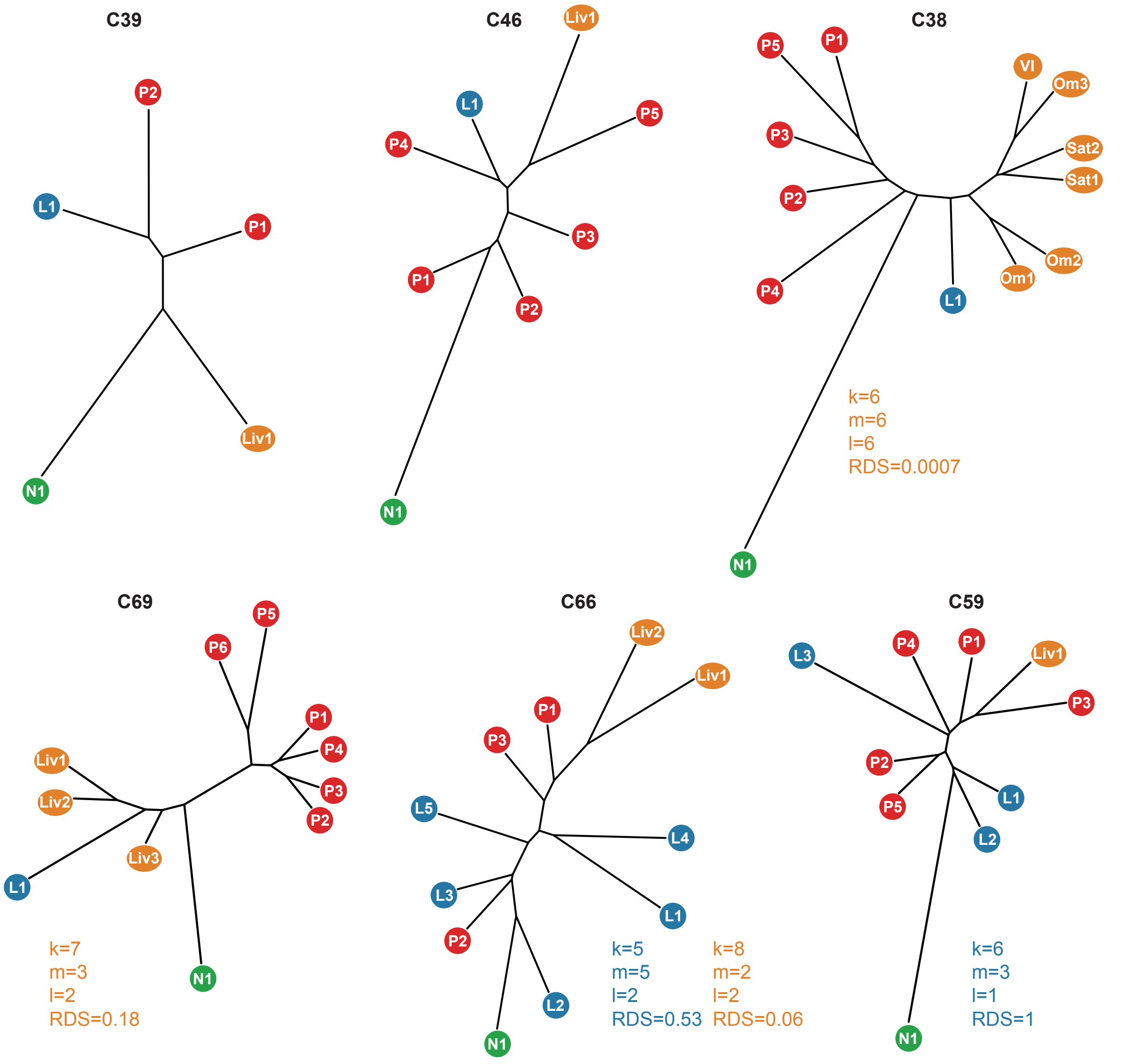


**Supplementary Fig. 2: Phylogenetic trees of colorectal cancer patients C39, C46, C38, C69, C66, and C59.** Trees adapted from Naxerova et al.[16]. N, normal tissue; P, primary tumor; L, lymph node metastasis; Liv, liver metastasis; Om, omentum; Sat, intra-colonic satellite nodule; VI, venous invasion; RDS, root diversity score. The RDS of lymphatic metastases is denoted in blue. The RDS of distant metastases is denoted in orange.

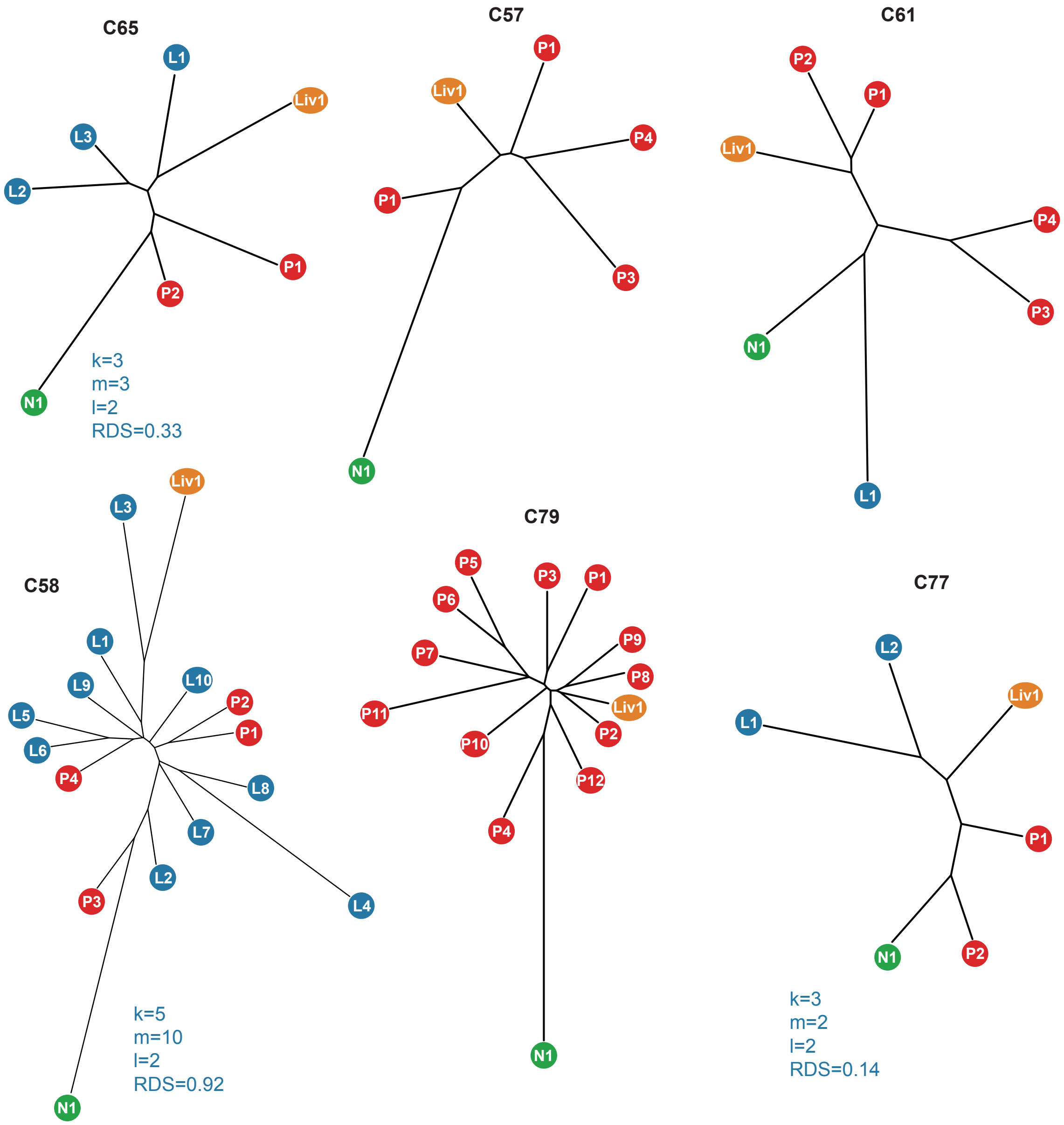


**Supplementary Fig. 3: Phylogenetic trees of colorectal cancer patients C65, C57, C61, C58, C79, and C77.** Trees adapted from Naxerova et al.[16]. N, normal tissue; P, primary tumor; L, lymph node metastasis; Liv, liver metastasis; RDS, root diversity score. The RDS of lymphatic metastases is denoted in blue.

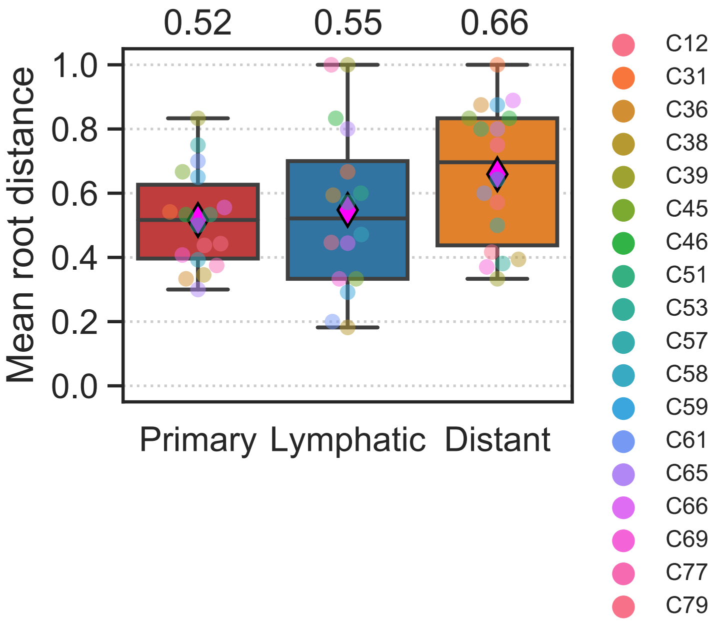


**Supplementary Fig. 4: Mean distances between the root normal sample and samples of primary tumors, lymphatic metastases, and distant metastases, respectively.** Distance was measured as the number of internal nodes separating a pair of samples and then normalized by the total number of internal nodes in a given phylogeny. Numbers on top denote the mean root distance in each group. Magenta diamond illustrates the mean in each group.

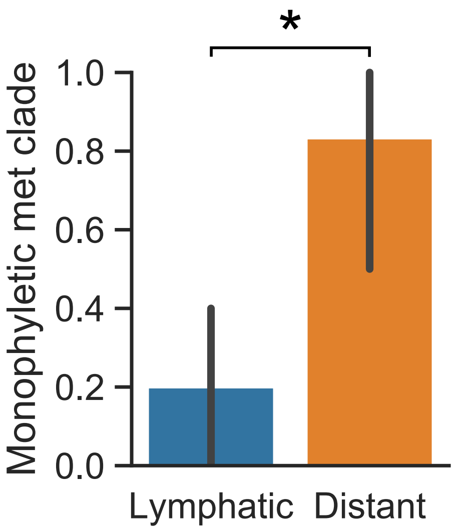


**Supplementary Fig. 5: Fraction of patients with a monophyletic clade of lymphatic or distant metastases.** According to a different classification scheme than in Fig. 1b, 20% (2/10) and 83% (5/6) of patients have one common ancestor for all lymphatic and distant metastases, respectively ($P = 0.035$, two-tailed Fishers exact test). Lymphatic or distant metastases were considered as a monophyletic group if the clade contained all lymphatic or distant metastases and no primary tumor samples. Samples of other metastases types can be part of the monophyletic group. See Supplementary Table 2 for individual classifications.

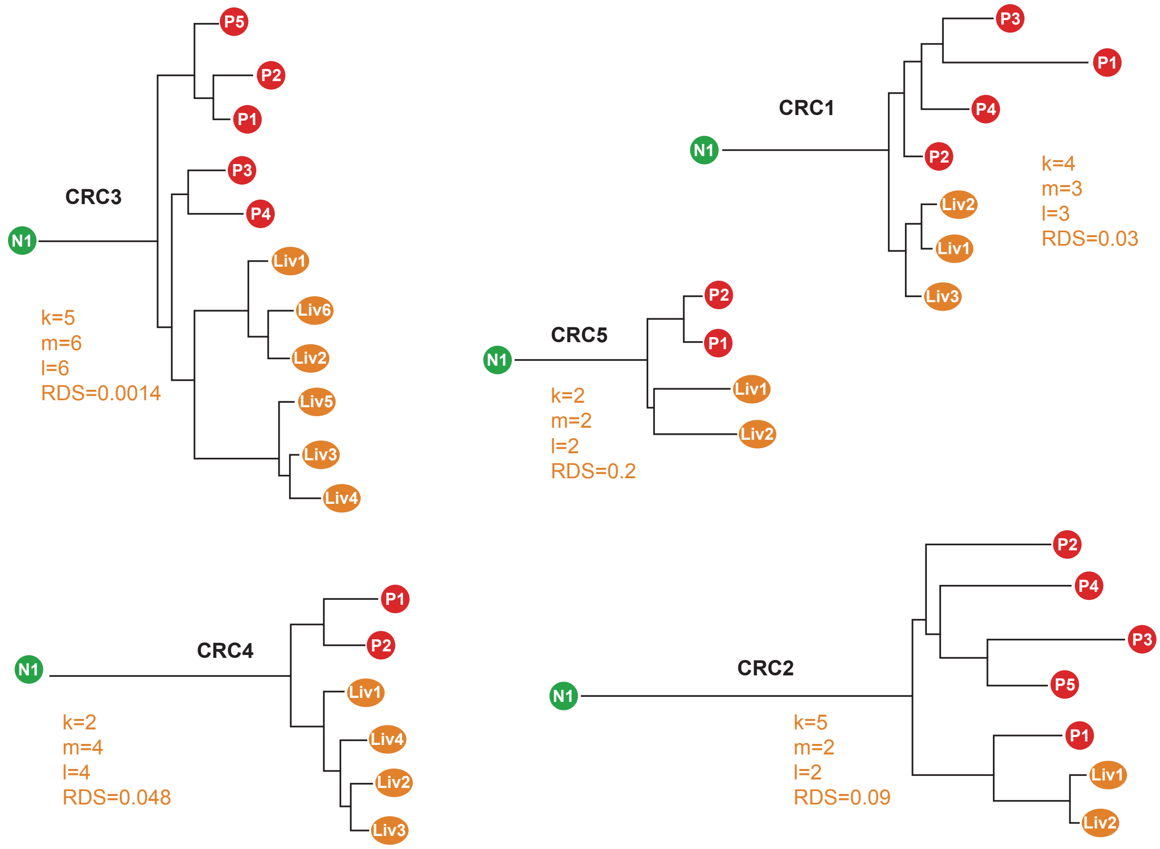


**Supplementary Fig. 6: Phylogenetic trees of colorectal cancer patients CRC1, CRC2, CRC3, CRC4, and CRC5.** Trees adapted from Kim et al.[18]. The RDS of distant metastases is denoted in orange. N, normal tissue; P, primary tumor; Liv, liver metastasis; RDS, root diversity score.

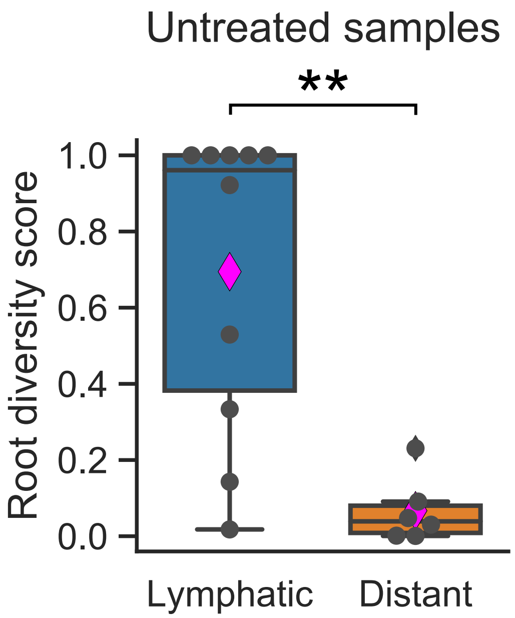


**Supplementary Fig. 7: Root diversity scores for untreated patients.** The root diversity score was significantly lower for distant metastases compared to lymphatic metastases (mean of 0.067 vs 0.69; $P = 7.0e\text{-}3$; two tailed Mann-Whitney test) in phylogenies from only untreated samples. Magenta diamond illustrates the mean root diversity score in each group. See Supplementary Table 2 for individual root diversity score values.

### Supplementary Methods: Root Diversity Score

Here we explain the *Root Diversity Score* and provide the mathematical intuition for the inferred formulas. We expand Edwards and Cavalli-Sforza's closed-form formula[28,29] that determines the number of bifurcating trees with $n$ external nodes: $\frac{(2n-3)!}{2^{n-2}(n-2)!}$. We denote the number of sequenced samples of distinct metastases of a given type (e.g., lymphatic or distant) as $m$ and the remaining number of sequenced cancer samples (not from this type of metastases) as $k$ per subject ($k, m \in \mathbb{N}$). The total number of analyzed cancer samples per subject is given by $k + m = n$. We describe a bifurcating tree as being of type $(k, m)$ if the samples from $m$ distinct metastases form one clade which does not consist of any other cancer samples. In other words, there is a single common ancestor of all metastases samples and this common ancestor does not contain any other samples. We refer to this type of clustering of metastases as a metastases clade. The total number of distinct tree topologies of type $(k, m)$ is denoted as $|(k, m)|$. A similar idea was used by Zhao et al. to calculate the probability that a single primary tumor sample would form an outgroup in an inferred phylogeny[26]. The root diversity score generalizes this approach to multiple sequenced primary tumor samples.

In Section 1, we present a recursive model that determines the number of bifurcating trees of type $(k, m)$. We infer a closed-form solution to calculate the number of bifurcating trees of type $(k, m)$.

In Section 2, we expand on the formulas in Section 1 by providing two recursive explanations for the number of trees of type $(k, m_1, m_2)$. A tree is described as type $(k, m_1, m_2)$ if there are $m_1 + m_2$ sequenced samples of distinct metastases which form exactly two metastases clades, one that contains $m_1$ samples and one that contains $m_2$ samples. We infer a closed-form equation for $|(k, m_1, m_2, ..., m_i)|$ where $m_1 + m_2 + ... + m_i$ is the total number of sequenced metastases.

In Section 3, we combine our results to calculate the probability that a tree with a metastases clade of a given size evolves by chance. The root diversity score can be calculated by using Equation (S2).

#### 1 Evolutionary trees with single root of metastases

To delineate the recursive relationship between the number of trees of type $(k + 1, m)$ and the number of trees of type $(k, m)$, we consider all 3-tip bifurcating trees. In total, there are three distinct topologies for $k = 1$ and $m = 2$ which are illustrated in Suppl. Fig. 8. Only one of these three trees is of type $(1, 2)$ because in the two other trees the metastases do not form a joint clade. Hence, $|(1, 2)| = 1$.

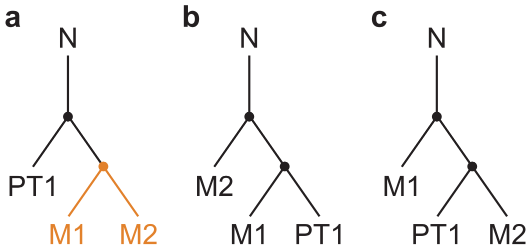


**Supplementary Fig. 8: All 3-tip bifurcating trees.** Metastases samples M1 and M2 form a clade (denoted in orange) only in tree illustrated in panel **a**. This tree is of type $(1, 2)$. N denotes the root of the tree (normal sample). PT1 denotes a sample of the primary tumor.

To obtain the number of trees where two metastases ($m = 2$) form a joint clade when $k = 2$ $|(2, 2)|$, we grow the tree of type $(1, 2)$ into all possible trees of type $(2, 2)$. Observe that in the type $(1, 2)$ tree (Suppl. Fig. 9a), there are exactly three (depicted in blue) out of $2 \cdot 3 - 1 = 5$ edges where we an additional sample PT2 (second sample of the primary tumor) can be added without dividing the metastases clade $\{$M1, M2$\}$. By adding sample PT2 to any of these three blue edges, three different 4-tip bifurcating trees of type $(2, 2)$ can form (Suppl. Fig. 9b). Thus, we claim that $|(2, 2)| = 3$.

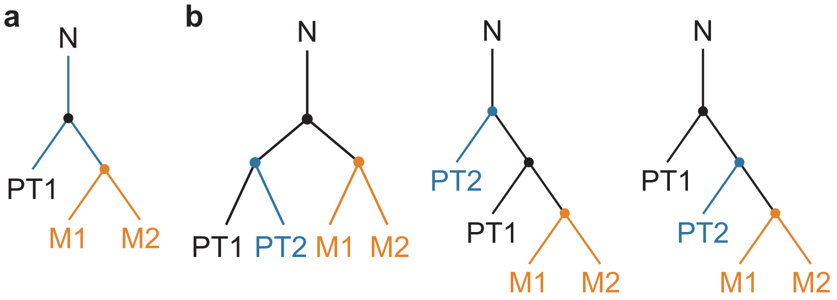


**Supplementary Fig. 9: Grow type $(2, 2)$ trees from type $(1, 2)$ trees. a** | There is only one tree of type $(1, 2)$ which can be grown into a tree of type $(2, 2)$ in three different ways. The three branches where the additional sample can be added to obtain a type $(2, 2)$ tree are depicted in blue. All other possible 4-tip trees would divide the metastases clade and hence a type $(3, 1)$ tree would arise. **b** | Three distinct type $(2, 2)$ trees exist which are inferred from extending the type $(1, 2)$ tree as illustrated in panel **a**. N denotes the root of the tree (normal sample). M denotes metastases samples and PT denotes primary tumor sample.

#### 1.1 Recursive formulas

The first theorem exemplifies the recursive relationship between the number of trees of type $(k+1, m)$ and the number of trees of type $(k, m)$.

**Lemma 1.** *Let $k, m \in \mathbb{N}$ such that $k > 1$. For every tree of type $(k, m)$, there exists a tree of type $(k-1, m)$ from which a type $(k, m)$ tree can be grown.*

*Proof.* Let $T$ be a tree of type $(k, m)$ where $k, m \in \mathbb{N}$ such that $k > 1$. For the sake of contradiction suppose that no tree of type $(k-1, m)$ exists from which $T$ can be grown. This would imply that $T$ is not reducible to any tree of type $(k-1, m)$. By definition of a tree of type $(k-1, m)$, this is a contradiction, since removing a non-metastatic sample from $T$ would result in a tree of type $(k-1, m)$. □

A similar proof can be created to illustrate analogous Lemma 2.

**Lemma 2.** *Let $k, m \in \mathbb{N}$ such that $m > 1$. For every tree of type $(k, m)$, there exists a tree of type $(k, m-1)$ from which a type $(k, m)$ tree can be grown.*

**Theorem 1.** *The number of type $(k+1, m)$ trees equals the number of type $(k, m)$ trees multiplied by $(2k+1)$: $|(k+1, m)| = |(k, m)| \cdot (2k+1)$.*

*Proof.* Let $m \in \mathbb{N}$ be fixed such that $m > 1$. We proceed via induction on $k$. First, consider the base case when $k = 1$. We aim to show that:

$$|(1+1, m)| = |(1, m)| \cdot (2 \cdot 1 + 1) \ .$$

Following Edwards and Cavalli-Sforza[28], every tree of type $(1, m)$ has $2 \cdot (m+1) - 1$ edges. Moreover, there are exactly $2 \cdot (m-1)$ edges on each tree of type $(1, m)$ where an additional sample cannot be added without dividing the metastases cluster of size $m$. Thus, there are

$$[2 \cdot (1+m) - 1] - [(m-1) \cdot 2] = [2 + 2m - 1] - [2m - 2] = 1 + 2 = 3$$

edges where the additional sample for every tree of type $(1, m)$ can be added. This implies,

$$|(1+1, m)| = |(1, m)| \cdot 3 = |(1, m)| \cdot (2 \cdot 1 + 1)$$

as desired. For the induction step, we assume

$$|(k+1, m)| = |(k, m)| \cdot (2 \cdot k + 1)$$

for some $k \in \mathbb{N}$ and we aim to show

$$|((k+1)+1, m)| = |(k+1, m)| \cdot (2(k+1)+1) \ .$$

Every tree of type $(k+1, m)$ has $2 \cdot (k+1+m) - 1 = 2k + 2m + 1$ edges. For each tree of type $(k+1, m)$, there exist $2 \cdot (m-1) = 2m - 2$ edges where an additional sample cannot be added without dividing the metastases cluster of size $m$. Hence, there are

$$(2k + 2m + 1) - (2m - 2) = 2k + 3$$

edges where the additional sample for each tree of type $(k+1, m)$ can be added. Thus,

$$|((k+1)+1, m)| = |(k+2, m)| = |(k+1, m)| \cdot (2k+3) = |(k+1, m)| \cdot (2(k+1)+1)$$

as desired. □

**Theorem 2.** *The number of trees where $m+1$ metastases form a clade is given by $|(k, m+1)| = |(k, m)| \cdot (2m-1)$.*

*Proof.* Let $k \in \mathbb{N}$ be fixed. We proceed via induction on $m$. First, we consider the base case where $m = 1$. We aim to show that:

$$|(k, 1+1)| = |(k, 1)| \cdot (2 \cdot 1 - 1) \ .$$

Every tree of type $(k, 1)$ has $2 \cdot (k+1) - 1$ edges. There are $2k$ edges out of all $2 \cdot (k+1) - 1$ edges where adding an additional metastasis sample would result in two separate metastases clades of size 1. Thus, there is

$$2 \cdot (k+1) - 1 - 2k = 1$$

edge on each tree of type $(k, 1)$ where an additional metastasis sample can be added such that it forms a clade with the already existing metastasis samples. This implies,

$$|(k, 1+1)| = |(k, 1)| \cdot 1 = |(k, 1)| \cdot (2 \cdot 1 - 1)$$

as desired.

For the induction step, we assume

$$|(k, m+1)| = |(k, m)| \cdot (2m-1)$$

and aim to show that

$$|(k, (m+1)+1)| = |(k, m+1)| \cdot (2(m+1)-1) \ .$$

Since every tree of type $(k, m+1)$ has $2 \cdot (k+m+1) - 1 = 2k+2m+1$ edges, there are $2k$ edges where adding an additional metastasis sample would result in two separate metastases clades. Thus, there are

$$(2k+2m+1) - (2k) = 2m+1$$

edges for each tree of type $(k, m+1)$ where the $(k+1)$th metastasis sample can be added such that it forms a clade with the already existing metastasis samples. This implies that

$$|(k, (m+1)+1)| = |(k, m+1)| \cdot (2m+1) = |(k, m+1)| \cdot (2(m+1)-1)$$

as desired.

□

#### 1.2 Closed-form solution

Next, we provide a closed-form formula to calculate the number of trees of type $(k, m)$.

**Theorem 3.** *For $m > 1$, the number of type $(k, m)$ trees is $|(k, m)| = \frac{(2k-1)!}{2^{k-1}(k-1)!} \cdot \frac{(2m-3)!}{2^{m-2}(m-2)!}$.*

To prove Theorem 3, we first consider Lemma 3.

**Lemma 3.** *The number of type $(k, 2)$ trees is given by $|(k, 2)| = \frac{(2k-1)!}{2^{k-1}(k-1)!}$.*

*Proof.* We start via induction on $k$. First, we consider the base case where $k = 1$. We aim to show that

$$|(1, 2)| = \frac{(2(1)-1)!}{2^{1-1}(1-1)!} \ .$$

Recall that the number of type $(1, 2)$ trees is $|(1, 2)| = 1$ (Suppl. Fig. 8a). We find that

$$|(1, 2)| = 1 = \frac{(2(1)-1)!}{2^{1-1}(1-1)!} \ .$$

For the induction step, we assume

$$|(k,2)| = \frac{(2k-1)!}{2^{k-1}(k-1)!}$$

for some $k \in \mathbb{N}$ and aim to show

$$|(k+1,2)| = \frac{(2(k+1)-1)!}{2^{(k+1)-1}((k+1)-1)!} \ .$$

Recall that by Theorem 1,

$$|(k+1,2)| = |(k,2)| \cdot (2k+1) \ .$$

By the induction hypothesis, we obtain

$$\begin{aligned}
|(k+1,2)| &= \frac{(2k-1)!}{2^{k-1}(k-1)!} \cdot (2k+1) \\
&= \frac{(2k+1)(2k-1)(2k-2)...(k)(k-1)...(1)}{2^k \cdot 2^{-1}(k-1)(k-2)...(1)} \\
&= \frac{(2k+1)(2k)(2k-1)(2k-2)...(k+1)}{2^k} \\
&= \frac{(2k+1)!}{2^k \cdot k!} \\
&= \frac{(2(k+1)-1)!}{2^{(k+1)-1}((k+1)-1)!}
\end{aligned}$$

as desired. □

Now we can prove Theorem 3.

*Proof.* Let $k \in \mathbb{N}$ be fixed. We proceed via induction on $m$. First, we consider the base case $m = 2$ and aim to show that

$$|(k,2)| = \frac{(2k-1)!}{2^{k-1}(k-1)!} \cdot \frac{(2(2)-3)!}{2^{(2)-2}((2)-2)!} \ .$$

Note that by Lemma 3

$$
\begin{aligned}
|(k,2)| &= \frac{(2k-1)!}{2^{k-1}(k-1)!} \\
&= \frac{(2k-1)!}{2^{k-1}(k-1)!}\cdot 1 \\
&= \frac{(2k-1)!}{2^{k-1}(k-1)!}\cdot \frac{(2(2)-3)!}{2^{(2)-2}((2)-2)!}\ .
\end{aligned}
$$

For the induction step, we assume

$$|(k,m)| = \frac{(2k-1)!}{2^{k-1}(k-1)!}\cdot\frac{(2m-3)!}{2^{m-2}(m-2)!}$$

for some $m > 1$ and aim to show

$$|(k,m+1)| = \frac{(2k-1)!}{2^{k-1}(k-1)!}\cdot\frac{(2(m+1)-3)!}{2^{(m+1)-2}((m+1)-2)!}\ .$$

Using Theorem 2, we obtain that

$$
\begin{aligned}
|(k,m+1)| &= |(k,m)|\cdot 2(m)-1 \\
&= \frac{(2k-1)!}{2^{k-1}(k-1)!}\cdot\frac{(2m-3)!}{2^{m-2}(m-2)!}\cdot 2(m)-1 \\
&= \frac{(2k-1)!}{2^{k-1}(k-1)!}\cdot\frac{(2m-1)(2m-3)(2m-4)...(m)(m-1)(m-2)...(1)}{2^m\cdot 2^{-2}(m-2)(m-3)...(1)} \\
&= \frac{(2k-1)!}{2^{k-1}(k-1)!}\cdot\frac{(2)(2m-1)(2m-2)(2m-3)(2m-4)...(m)}{2^m} \\
&= \frac{(2k-1)!}{2^{k-1}(k-1)!}\cdot\frac{(2m-1)!}{2^m 2^{-1}(m-1)!} \\
&= \frac{(2k-1)!}{2^{k-1}(k-1)!}\cdot\frac{(2(m+1)-3)!}{2^{(m+1)-2}((m+1)-2)!}
\end{aligned}
$$

as desired. □

### 2 Evolutionary trees with multiple roots of metastases

A tree is described as type $(k, m_1, m_2)$ if it has $m_1 + m_2$ sampled metastases such that there are two distinct metastases clades of size $m_1$ and $m_2$ as well as $k$ other samples. In this section, we provide the intuition for calculating the number of type $(k, m_1, m_2)$ trees $|(k, m_1, m_2)|$.

We first consider all trees of type $(1, 1, 1)$ (Suppl. Fig. 10). There are a total of three 3-tip bifurcating trees. Each of these trees is of type $(1, 1, 1)$ since we now assume that each metastasis forms a clade of its own kind of size 1, hence $|(1, 1, 1)| = 3$ (compare Figs. 8 and 10).

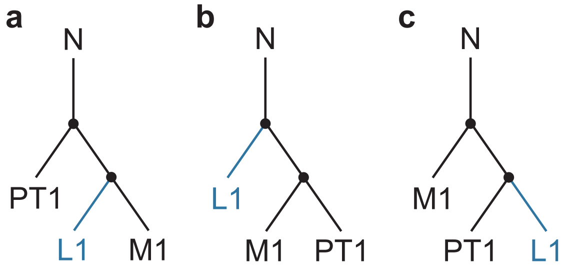


**Supplementary Fig. 10: All 3-tip bifurcating trees for three classes of samples.** Metastases clades of two kinds (M and L) exist. To grow the metastases L kind clade, an additional sample could be added at the blue edges. M denotes distant metastases sample, L lymphatic metastases sample, and PT denotes primary tumor sample. N denotes the root of the tree (normal sample).

To infer the number of type $(1, 1, 2)$ trees $|(1, 1, 2)|$, we consider all trees of type $(1, 1, 2)$ that can be grown from the three trees of type $(1, 1, 1)$ (see blue branches in Suppl. Fig. 10). In each of the three trees of type $(1, 1, 1)$, there is $2 \cdot m_2 - 1 = 2 \cdot 1 - 1 = 1$ edge (the blue edge) where an additional sample L2 could be added such that $m_2 = 2$ and a tree of type $(1, 1, 2)$ is formed (Suppl. Fig. 11). We claim that these are the only trees of type $(1, 1, 2)$, or equivalently, all trees of type $(1, 1, 2)$ can be grown from the three trees of type $(1, 1, 1)$. This result is an extension of Lemma 2. Thus, the number of type $(1, 1, 2)$ trees is $|(1, 1, 2)| = |(1, 1, 1)| \cdot 1 = 3$.

#### 2.1 Recursive formulas

We present recursive formulas for type $(k, m_1, m_2)$ trees when there are two different kinds of metastases forming clades with $m_1$ and $m_2$ samples, respectively.

**Theorem 4.** *The number of type* $(k, m_1 + 1, m_2)$ *trees is given by* $|(k, m_1 + 1, m_2)| = |(k, m_1, m_2)| \cdot (2m_1 - 1)$ *for all* $m_1 \in \mathbb{N}$.

*Proof.* Without loss of generality, we fix $k, m_2 \in \mathbb{N}$. We proceed via induction on $m_1$. We

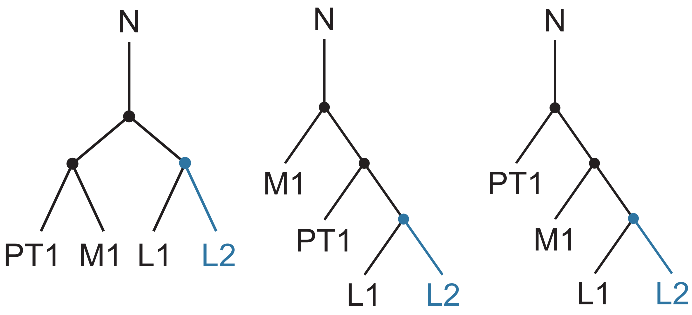


**Supplementary Fig. 11: All trees of type (1,1,2).** Two metastases of kind L form a clade as well as metastasis M1 forms its own clade. M denotes distant metastases sample, L lymphatic metastases sample, and PT denotes primary tumor sample. N denotes the root of the tree (normal sample).

start with the base case where $m_1 = 1$ and aim to show that

$$|(k, 1+1, m_2)| = |(k, 1, m_2)| \cdot (2(1) - 1) \ .$$

Since every tree of type $(k, 1, m_2)$ has $2 \cdot (k + 1 + m_2) - 1$ edges, there are $2(k + m_2)$ edges for each tree of type $(k, 1, m_2)$ where we cannot add the second metastasis of the first kind without forming a separate clade. Thus, there is

$$[2(k + 1 + m_2) - 1] - [2(k + m_2)] = 1$$

edge on each tree of type $(k, 1, m_2)$ where we can add a second metastasis of the first kind such that it forms a clade with the existing one. This implies,

$$|(k, 1+1, m_2)| = |(k, 1, m_2)| \cdot 1 = |(k, 1, m_2)| \cdot (2(1) - 1)$$

as desired. For the induction step, we assume

$$|(k, m_1 + 1, m_2)| = |(k, m_1, m_2)| \cdot (2m_1 - 1)$$

and aim to show that

$$|(k, (m_1 + 1) + 1, m_2)| = |(k, m_1 + 1, m_2)| \cdot (2(m_1 + 1) - 1) \ .$$

Every tree of type $(k, m_1 + 1, m_2)$ has $2 \cdot (k + m_1 + 1 + m_2) - 1 = 2k + 2m_1 + 2 + 2m_2$ edges. Because there are $2(k + m_2)$ edges where the $(m_1 + 2)$th metastasis of the first kind cannot be added without forming a separate cluster, there are

$$(2k + 2m_1 + 2 + 2m_2) - (2k + 2m_2) = 2m_1 + 2$$

edges for each tree of type $(k, m + 1, m_2)$ where the $(m_1 + 2)$th metastasis of the first kind could be added. This implies that

$$|(k, (m_1 + 1) + 1, m_2)| = |(k, m_1 + 1, m_2)| \cdot (2m_1 + 2) = |(k, m_1 + 1, m_2)| \cdot (2(m_1 + 1) - 1)$$

as desired. □

A similar proof can be created to prove Theorem 5.

**Theorem 5.** *The number of type $(k, m_1, m_2 + 1)$ trees is given by $|(k, m_1, m_2 + 1)| = |(k, m_1, m_2)| \cdot (2m_2 - 1)$ for all $m_2 \in \mathbb{N}$.*

**Theorem 6.** *The number of type $(k + 1, m_1, m_2)$ trees is given by $|(k + 1, m_1, m_2)| = |(k, m_1, m_2)| \cdot (2k + 3)$ for all $k \in \mathbb{N}$.*

*Proof.* Without loss of generality, we fix $m_1, m_2 \in \mathbb{N}$. We proceed via induction on $k$. We consider the base case where $k = 1$ and aim to show that

$$|(1 + 1, m_1, m_2)| = |(1, m_1, m_2)| \cdot (2(1) + 3) \ .$$

Every tree of type $(1, m_1, m_2)$ has $2 \cdot (m_1 + m_2 + 1) - 1$ edges. There are $2 \cdot (m_1 - 1 + m_2 - 1)$ edges on each tree of type $(1, m_1, m_2)$ where a second sample cannot be added without dividing one of the distinct metastasis clades, $m_1$ or $m_2$. Thus, there are

$$[2 \cdot (m_1 + m_2 + 1) - 1] - 2 \cdot (m_1 - 1) - 2 \cdot (m_2 - 1) = (2m_1 + 2m_2 + 2 - 1) - 2m_1 + 2 - 2m_2 + 2 = 1 + 2 + 2 = 5$$

edges where the second sample for every tree of type $(1, m_1, m_2)$ can be added without forming a third metastasis clade. This implies,

$$|(1 + 1, m_1, m_2)| = |(1, m_1, m_2)| \cdot 5 = |(1, m_1, m_2)| \cdot (2(1) + 3)$$

as desired. For the induction step, we assume

$$|(k + 1, m_1, m_2)| = |(k, m_1, m_2)| \cdot (2(k) + 3)$$

for some $k \in \mathbb{N}$ and we aim to show that

$$|((k+1)+1, m_1, m_2)| = |(k+1, m_1, m_2)| \cdot (2(k+1)+3) \ .$$

Since every tree of type $(k+1, m_1, m_2)$ has $2 \cdot (k+1+m_1+m_2) - 1 = 2k+2m_1+2m_2+1$ edges, there are $2 \cdot (m_1+m_2-2)$ edges where the $k+1$th sample cannot be added without dividing at least one of the distinct metastases clades, $m_1$ or $m_2$. Thus, there are

$$(2k+2m_1+2m_2+1) - (2m_1+2m_2-4) = 2k+5$$

edges for each tree of type $(k+1, m_1, m_2)$ where the $k+1$th sample can be added such that the metastases clades remain intact. Thus,

$$|((k+1)+1, m_1, m_2)| = |(k+2, m_1, m_2)| = |(k+1, m_1, m_2)| \cdot (2k+5) = |(k+1, m_1, m_2)| \cdot (2(k+1)+3)$$

as desired. □

#### 2.2 Closed-form formulas

We present a closed-form solution to calculate the number of type $(k, m_1, m_2, ...., m_i)$ trees for any number of different metastases kinds $i \in \mathbb{N}$ that each form its own clade.

**Theorem 7.** *The number of type $(k, m_1, m_2, ...., m_i)$ trees for any $i \in \mathbb{N}$ different metastases kinds that each form its own clade is*

$$|(k, m_1, m_2, ..., m_i)| = \left( \frac{(2k+(2i-3))!}{2^{k+(i-2)}(k+(i-2))!} \right) \left( \prod_{j=1}^{i} \frac{(2m_j-3)!}{2^{m_j-2}(m_j-2)!} \right) .$$

To prove Theorem 7, we use the following Lemma 4.

**Lemma 4.** *The number of type $(k, m_1, m_2, ..., m_i, m_{i+1})$ for any given $i \in \mathbb{N}$ is given by*

$$|(k, m_1, m_2, ..., m_i, m_{i+1})| = |(k, m_1, m_2, ..., m_i)|(2k+2i-1) \left( \prod_{j=1}^{m_{i+1}-1} 2(j) - 1 \right)$$

*for all $m_{i+1} \in \mathbb{N}$.*

*Proof.* Let $k, m_1, m_2, ..., m_i \in \mathbb{N}$ for some $i \in \mathbb{N}$. We proceed via induction on $m_{i+1}$. We first consider the base case $m_{i+1} = 1$ and aim to show that

$$|(k, m_1, m_2, ..., m_i, 1)| = |(k, m_1, m_2, ..., m_i)|(2k + 2i - 1) \ .$$

An arbitrary tree $T$ of type $(k, m_1, m_2, ..., m_i)$ has $2(k+m_1+...+m_i)-1$ edges. Moreover, $T$ has $2(m_1+...+m_i-i)$ edges where we cannot add a sample of a new metastasis kind without separating one of the existing metastases clades. Thus, there are $(2k+2m_1+...+2m_i-1)-(2m_1+...+2m_i-2i) = 2k+2i-1$ edges where we can place the $(k+m_1+m_2+...+m_i+m_{i+1})th$ metastasis. Since $T$ was an arbitrary chosen tree of type $(k, m_1, m_2, ..., m_i)$,

$$|(k, m_1, m_2, ..., m_i, 1)| = |(k, m_1, m_2, ..., m_i)|(2k + 2i - 1)$$

as desired.

For the induction step, we assume

$$|(k, m_1, m_2, ..., m_i, m_{i+1})| = |(k, m_1, m_2, ..., m_i)|(2k + 2i - 1)\left(\prod_{j=1}^{m_{i+1}-1} 2j - 1\right)$$

for some $m_{i+1} \in \mathbb{N}$ and aim to show that

$$|(k, m_1, m_2, ..., m_i, m_{i+1} + 1)| = |(k, m_1, m_2, ..., m_i)|(2k + 2i - 1)\left(\prod_{j=1}^{m_{i+1}} 2j - 1\right) \ .$$

We consider the right hand side of the above equation. By our induction hypothesis and Theorem 4, we have

$$\begin{aligned} &|(k, m_1, m_2, ..., m_i)|(2k + 2i - 1)\left(\prod_{j=1}^{m_{i+1}} 2j - 1\right) \\ &= |(k, m_1, m_2, ..., m_i)|(2k + 2i - 1)\left(\prod_{j=1}^{m_{i+1}-1} 2j - 1\right)(2m_{i+1} - 1) \\ &= |(k, m_1, m_2, ..., m_i, m_{i+1})|(2m_{i+1} - 1) \\ &= |(k, m_1, m_2, ..., m_i, m_{i+1} + 1)| \end{aligned}$$

as desired. □

Next we present the proof for Theorem 7.

*Proof.* Let $k, m_t \in \mathbb{N}$ for all $t \in \mathbb{N}$. We proceed via induction on $i$. We first consider the base case $i = 1$ and aim to show that

$$|(k, m_1)| = \left(\frac{[2k + (2(1) - 3)]!}{2^{k+(1-2)}[k + (1-2)]!}\right)\left(\prod_{j=1}^{1} \frac{(2m_j - 3)!}{2^{m_j-2}(m_j - 2)!}\right).$$

By algebraic simplification and Theorem 3 we know that

$$\left(\frac{[2k + (2(1) - 3)]!}{2^{k+(1-2)}[k + (1-2)]!}\right)\left(\prod_{j=1}^{1} \frac{(2m_j - 3)!}{2^{m_j-2}(m_j - 2)!}\right) = \frac{(2k-1)!}{2^{k-1}(k-1)!} \cdot \frac{(2m_1 - 3)!}{2^{m_1-2}(m_1 - 2)!} = |(k, m_1)|$$

as desired.

For the induction step, we assume that

$$|(k, m_1, m_2, ..., m_i)| = \left(\frac{(2k + (2i - 3))!}{2^{k+(i-2)}(k + (i-2))!}\right)\left(\prod_{j=1}^{i} \frac{(2m_j - 3)!}{2^{m_j-2}(m_j - 2)!}\right)$$

for some $i \in \mathbb{N}$. We aim to show that

$$|(k, m_1, m_2, ..., m_i, m_{i+1})| = \left(\frac{[2k + (2(i+1) - 3)]!}{2^{k+((i+1)-2)}[k + ((i+1) - 2)]!}\right)\left(\prod_{j=1}^{i+1} \frac{(2m_j - 3)!}{2^{m_j-2}(m_j - 2)!}\right).$$

Again by algebraic simplification, the induction hypothesis, and Lemma 4, we know that

$$\left(\frac{[2k + (2(i+1) - 3)]!}{2^{k+((i+1)-2)}[k + ((i+1) - 2)]!}\right)\left(\prod_{j=1}^{i+1} \frac{(2m_j - 3)!}{2^{m_j-2}(m_j - 2)!}\right) =$$

$$= \left(\frac{[2k + 2i - 1]!}{2^{k+i-1}(k + i - 1)!}\right)\left(\prod_{j=1}^{i+1} \frac{(2m_j - 3)!}{2^{m_j-2}(m_j - 2)!}\right)$$

$$= \left(\frac{[2k + 2i - 3]!}{2^{k+i-2}(k + i - 2)!}\right)\left(\prod_{j=1}^{i} \frac{(2m_j - 3)!}{2^{m_j-2}(m_j - 2)!}\right)\left(\frac{(2k + 2i - 2)(2k + 2i - 1)}{2(k + i - 1)}\right)\left(\frac{(2m_{i+1} - 3)}{2^{m_{i+1}-2}(m_{i+1} - 2)}\right)$$

$$= |(k, m_1, m_2, ..., m_i)|\left(\frac{(2k + 2i - 2)(2k + 2i - 1)}{2k + 2i - 2}\right)\left(\frac{(2m_{i+1} - 3)}{2^{m_{i+1}-2}(m_{i+1} - 2)}\right)$$

$$= |(k, m_1, m_2, ..., m_i)|(2k + 2i - 1)\left(\frac{(2m_{i+1} - 3)}{2^{m_{i+1}-2}(m_{i+1} - 2)}\right)$$

$$= |(k, m_1, m_2, ..., m_i, m_{i+1})|$$

as desired. $\square$

### 3 Probability of clade existence

Now that we can calculate the number of different tree topologies where a given number of metastases cluster together and we also know the total number of distinct trees for a given number of samples[28], we can compute the probability that such a tree with a cluster of metastases arises by chance. Using the inclusion-exclusion principle, we can then calculate a *root Diversity Score* given by the probability that a tree with a metastases cluster of a certain size or larger would arise by chance. Note that while we focused here on quantifying the root diversity of distant metastases, this approach can equally be applied to clusters of any other type (e.g., lymphatic metastases or regions of the primary tumor; **Fig. 2**).

We calculate the probability that at least $l$ metastases form a clade ($1 \leq l \leq m$) given a bifurcating tree with $m$ metastases samples and $n = m + k$ samples in total. In other words, of all bifurcating trees with $n$ leaves, how many exhibit a clade of size $\{l, l+1, .., m\}$. Using the inclusion-exclusion principle, the number of $n$-bifurcating trees that exhibit a metastasis clade of size $\{l, l+1, .., m\}$ is given by

$$\sum_{i=1}^{\lfloor \frac{m}{l} \rfloor} (-1)^{i-1} \frac{\prod_{j=0}^{i-1} \binom{m-jl}{l}}{i!} |(n - i \times l, \{l\}^i)| \ . \tag{S1}$$

Then the probability that at least $l$ out of $m$ metastases form a clade in any $n$-bifurcating tree is given by

$$\Pr(c(T(k,m)) \geq l) = \frac{2^{n-2}(n-2)!}{(2n-3)!} \sum_{i=1}^{\lfloor \frac{m}{l} \rfloor} (-1)^{i-1} \frac{\prod_{j=0}^{i-1} \binom{m-j \cdot l}{l}}{i!} |(n - i \times l, \{l\}^i)| \tag{S2}$$

where $c(T(k,m))$ denotes the largest metastases clade in a bifurcating tree with $m$ metastases samples and $k+m$ samples in total. At `https://github.com/johannesreiter/rootdiversity`, an implementation of this framework is provided. (WILL BECOME AVAILABLE WITH ACCEPTANCE OF THIS MANUSCRIPT; FOR REVIEW PLEASE SEE SUPPLEMENTARY FILES).

To provide an intuition for Equation (S2) that calculates the root diversity score, we present the following example. Suppose we have an 11-tip bifurcating tree where $k = 5$, $m = 6$, and $l = 2$. We assume five primary tumor samples $\{PT1, PT2, PT3, PT4, PT5\}$

and samples from six distinct metastases $\{M1, M2, M3, M4, M5, M6\}$. We further assume that the inferred cancer phylogeny shows that only $\{M1, M2\}$ form a common clade. Hence, of all 11-tip bifurcating trees, we need to calculate how many of the bifurcating trees exhibit a cluster of sizes $2, 3, 4, 5$ or 6. Since each cluster of size $3, 4, 5$ or 6 also contains a cluster of size 2, we enumerate all 11-tip bifurcating trees that have a cluster of size 2. By Theorem 3, we know that the number of 11-tip bifurcating trees with $k = 5$ primary tumor samples and $m = 6$ metastases where two metastases form a clade, is given by $|(k+m-l, l)| = |(9, 2)|$. However, the cluster size of 2 may contain any two of the 6 metastasis:$\{M1, M2, M3, M4, M5, M6\}$. So the total number of distinct 11-tip bifurcating trees with a cluster size of 2 is $\binom{6}{2} \times |(9, 2)|$.

Because there are many tree topologies that exhibit multiple distinct metastases clades of size 2, we have to compensate for this overcounting by subtracting the number of trees that exhibit multiple clades. Hence, we subtract the number of 11-tip bifurcating trees with two distinct clusters of size 2 that have any combination of metastases $\{M1, M2, M3, M4, M5, M6\}$

$$\binom{6}{2}|(9,2)| - \frac{\binom{6}{2}\binom{4}{2}}{2!}|(4,2,2)| \ .$$

However, because some 11-tip bifurcating trees exhibit even three distinct metastases clades of size 2 and therefore have three different combinations with two clades of size, we now undercounted the number of 11-tip bifurcating trees that allow for a clustering size of $2, 3, 4, 5$ or 6. Again following the inclusion-exclusion principle, we add those trees with three distinct clusters of size 2 that have any combination of metastases $\{M1, M2, M3, M4, M5, M6\}$. Hence, we obtain for the number of 11-tip bifurcating trees that exhibit a clade of size $2, 3, 4, 5$, or 6:

$$\binom{6}{2}|(9,2)| - \frac{\binom{6}{2}\binom{4}{2}}{2!}|(7,2,2)| + \frac{\binom{6}{2}\binom{4}{2}\binom{2}{2}}{3!}|(5,2,2,2)| \ .$$

Thus, the probability that at least $l = 2$ metastases form a clade given $k = 5$ and $m = 6$ in a 11-tip bifurcating tree is

$$\frac{2^{8-2}(8-2)!}{(2(8)-3)!}\left[\binom{6}{2}|(6,2)| - \frac{\binom{6}{2}\binom{4}{2}}{2!}|(4,2,2)| + \frac{\binom{6}{2}\binom{4}{2}\binom{2}{2}}{3!}|(2,2,2,2)|\right] = 0.653 \ .$$
